# Origins of the current outbreak of multidrug resistant malaria in Southeast Asia: a retrospective genetic study

**DOI:** 10.1101/208371

**Authors:** Roberto Amato, Richard D. Pearson, Jacob Almagro-Garcia, Chanaki Amaratunga, Pharath Lim, Seila Suon, Sokunthea Sreng, Eleanor Drury, Jim Stalker, Olivo Miotto, Rick M. Fairhurst, Dominic P. Kwiatkowski

## Abstract

**Background:** Antimalarial failure is rapidly spreading across parts of Southeast Asia where dihydroartemisinin-piperaquine (DHA-PPQ) is used as first line treatment. The first published reports came from western Cambodia in 2013. Here we analyse genetic changes in the *Plasmodium falciparum* population of western Cambodia in the six years prior to that.

**Methods:** We analysed genome sequence data on 1492 *P. falciparum* samples from Southeast Asia, including 464 collected in western Cambodia between 2007 and 2013. Different epidemiological origins of resistance were identified by haplotypic analysis of the *kelch13* artemisinin resistance locus and the *plasmepsin 2-3* piperaquine resistance locus.

**Findings:** We identified over 30 independent origins of artemisinin resistance, of which the KEL1 lineage accounted for 91% of DHA-PPQ-resistant parasites. In 2008, KEL1 combined with PLA1, the major lineage associated with piperaquine resistance. By 2012, the KEL1/PLA1 co-lineage had reached over 60% frequency in western Cambodia and had spread to northern Cambodia.

**Interpretation:** The KEL1/PLA1 co-lineage emerged in the same year that DHA-PPQ became the first line antimalarial drug in western Cambodia and spread aggressively thereafter, displacing other artemisinin-resistant parasite lineages. These findings have significant implications for management of the global health risk associated with the current outbreak.

**Funding:** Wellcome Trust, Bill & Melinda Gates Foundation, Medical Research Council, UK Department for International Development, and Intramural Research Program of the US National Institute of Allergy and Infectious Diseases, National Institutes of Health.

## 1 Introduction

The first-line treatment for *Plasmodium falciparum* malaria is artemisinin combination therapy (ACT).^1^ Artemisinin and its derivatives are potent and fast-acting antimalarial drugs but they are also relatively short-acting.^2^ The rationale of ACT is to combine artemisinin with a longer-acting partner drug to ensure that all parasites are killed and also to prevent the emergence of resistance.^3^

In 2008 it became apparent that *P. falciparum* was becoming resistant to artemisinin in western Cambodia, and over the next few years the same phenomenon was observed in other parts of Cambodia as well as Thailand, Vietnam, Myanmar, and Laos.^4–6^ Although this caused widespread concerns there were some mitigating factors. First, the resistance to artemisinin was not complete, i.e. artemisinin treatment continued to reduce parasitaemia, albeit at a slower rate than previously. Second, ACT partner drugs were able to clear parasites despite their slower response to artemisinin, so that ACT remained effective as a first-line antimalarial therapy. Third, the spread of resistance was due to multiple emergences of artemisinin resistance, each caused by an independent mutation in the *kelch13* gene and confined to a relatively small geographical area.^7,8^

In 2013 it was reported that the situation was worsening, in that ACT was completely failing to clear parasites in some patients in western Cambodia.^9^ At that time, the form of ACT used in Cambodia as first-line treatment was dihydroartemisinin-piperaquine (DHA-PPQ) and it was found that ACT failure was due to increasing resistance to piperaquine. Piperaquine resistance was shown to have a genetic basis and copy number amplification of the *plasmepsin 2* and *plasmepsin 3* genes was discovered to be a useful genetic marker.^10,11^ Since 2013 the frequency of complete treatment failure in patients receiving the DHA-PPQ form of ACT has risen rapidly in Cambodia, northeast Thailand and Vietnam.^12–16^

It has been reported that this outbreak of DHA-PPQ treatment failure is associated with a specific parasite lineage, designated by the authors as *PfPailin*, that is spreading across the region.^14,16^ Although the parasites remain sensitive to other forms of ACT, these reports have caused considerable alarm and debate about whether or not it constitutes a public health emergency. We have conducted an in-depth genetic analysis of the epidemiological origins and early history of this outbreak, using genome sequence data on samples collected in Cambodia from 2007 to 2013, complemented by open access sequence data on other Southeast Asian samples generated by the MalariaGEN *Plasmodium falciparum* Community Project.^17^

## Results

### Multiple origins of *kelch13* resistance alleles

We analysed genome sequence data on 1492 samples collected at 11 locations across Southeast Asia, including 464 samples collected over seven consecutive years between 2007 and 2013 in Cambodia (Table 1). We began by studying non-synonymous single-nucleotide substitutions (SNPs) in the *P. falciparum kelch13* gene that are known or candidate markers of artemisinin resistance.^18^ We refer to these as *kelch13* resistance alleles. Overall, 46% of samples (n=689) carried a *kelch13* resistance allele; this includes 136 samples that were heterozygous due to mixed infection (Supplementary Table 1). We observed 24 distinct *kelch13* resistance alleles, each denoted by their effect on the amino-acid sequence, e.g. 538V (Supplementary Table 1, Figure 1). The most common resistance alleles were 580Y (accounting for 57% of resistant samples), 493H (10%), 539T (8%), 543T (4%), 441L (3%), 561H (3%), and 675V (3%).

**Table 1.** The 1492 samples analysed in this study, according to region and year.

| Region | Country/Site | 2002-2006 | 2007 | 2008 | 2009 | 2010 | 2011 | 2012 | 2013 | Total |
| --- | --- | --- | --- | --- | --- | --- | --- | --- | --- | --- |
| South Asia | Bangladesh |  |  | 9 | 11 |  |  | 34 |  | 54 |
| Southeast Asia<br>(West) | Myanmar |  |  |  |  |  | 56 | 44 | 1 | 101 |
|  | Thailand (Northwest, South) | 89 | 9 | 83 | 1 | 7 | 39 | 74 | 31 | 333 |
| Southeast Asia<br>(East) | Thailand (Northeast) |  |  |  |  |  | 4 | 12 | 3 | 19 |
|  | Laos |  |  |  |  | 33 | 40 | 23 |  | 96 |
|  | Vietnam |  |  |  | 9 | 57 | 88 | 23 |  | 177 |
|  | Cambodia (Northeast) |  |  |  |  | 36 | 59 | 20 | 10 | 125 |
|  | Cambodia (North) |  |  |  |  |  | 61 | 41 | 21 | 123 |
|  | Cambodia (West) |  | 25 | 41 | 58 | 102 | 141 | 59 | 38 | 464 |

**Figure 1.**
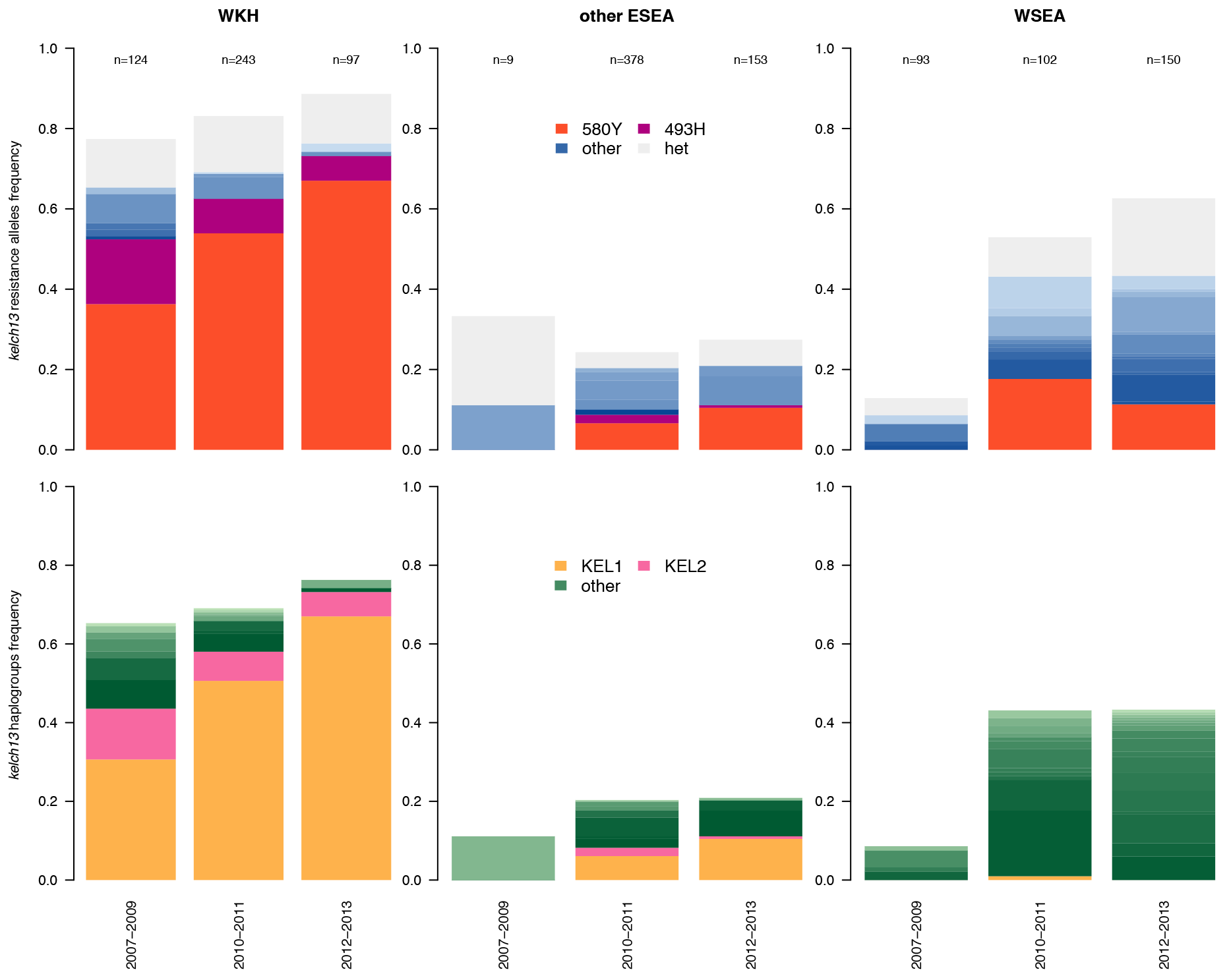
Figure 1. Frequency of *kelch13* resistance alleles (top) and haplogroups (bottom). Each panel reports the cumulative frequency of *kelch13* resistance alleles (top) and haplogroups (bottom) in three consecutive year groups in western Cambodia (left panels); northern and northeastern Cambodia, Vietnam, Laos, and northeastern Thailand (central panels); and northwestern and southern Thailand and Myanmar (right panels). In the top three panels, the frequency of the two most common *kelch13* resistance alleles are highlighted in red and purple (indicating 580Y and 493H, respectively) while all other resistance alleles are reported in shades of blue (homozygous) and grey (heterozygous). In the bottom three panels, the two most common *kelch13* haplogroups are indicated in yellow and pink (identifying KEL1 and KEL2, respectively), while all other haplogroups are reported in shades of green. The sample size for each group is reported at the top.

*Kelch13* resistance alleles were observed throughout the entire region, except in Bangladesh, and increased in frequency over time (Figure 1). Western Cambodia presented an extremely high prevalence of *kelch13* resistance alleles (83% overall, including 13% heterozygous) and their frequency steadily increased from 40% in 2007 to 84% in 2013. Of those samples with a homozygous *kelch13* resistance mutation in western Cambodia, 75% (n=241/323) were carrying the 580Y mutation (Supplementary Table 1).

The observation of 24 different resistance alleles indicates at least 24 epidemiological origins of artemisinin resistance in the sampled locations. However, some resistance alleles have multiple independent origins, so that two samples with the same resistance allele are not necessarily epidemiologically related, i.e. their alleles are identical by state but they might not be identical by descent.^7,8^ To address this question, we analysed the haplotype structure of *kelch13* and its flanking regions in all 553 samples with a homozygous resistance allele, and used statistical chromosome painting to estimate the level of shared ancestry between samples.^19^ This approach allowed us to assign each sample to a *kelch13* haplogroup, defined here as a group of samples with the same *kelch13* resistance mutation and strong haplotypic similarities indicative of recent shared lineage at the *kelch13* locus (Supplementary Figure 1). This approach identified 38 *kelch13* haplogroups, each of which is presumed to represent a distinct lineage of artemisinin resistance (Supplementary Table 2). The 580Y resistance allele was found in six different *kelch13* haplogroups, of which the most common was denoted the KEL1 lineage.

The KEL1 lineage accounts for 48% of artemisinin-resistant samples in this dataset. It is predominantly found in western Cambodia but is also observed in other parts of Cambodia as well as Vietnam and Laos, where it appears to have moved onto different genetic backgrounds (Figure 1, Supplementary Table 2 and Supplementary Figure 2). In western Cambodia, the KEL1 lineage was at 4% frequency in 2007, and rose to 63% frequency in 2013. It appears to have spread through the parasite population by recombination, i.e. it is found on parasites that are considerably different at the whole-genome level, as can be visualised with a genome-wide neighbour-joining tree (Supplementary Figure 2).

### Single major origin of *plasmepsin 2-3* amplifications

*Plasmepsin 2-3* amplifications, which are markers of piperaquine resistance, were observed in 41% (185/456) of samples from western Cambodia and in 1% (14/1009) of samples from elsewhere (Figure 2 and Supplementary Table 3; calls were inconclusive for 27 samples). Two genetic features of the *plasmepsin 2-3* amplifications indicated that they were mostly derived from the same epidemiological origin. First, the breakpoint sequences of the amplification, i.e. the sequences around the point where the additional copy of the gene is inserted, were identical in all but six samples (Supplementary Figure 3). Second, most samples carrying the amplification had highly similar haplotypes surrounding the amplified genes, which were different from those without the amplification (Supplementary Figure 3). Based on haplotype analysis, 93% (186/199) of samples with the amplification had strong evidence of recent shared ancestry (<5 differences out of 1454 SNPs with minor allele frequency >1%), and 6% (11/199) of samples could plausibly have arisen from the same recent ancestor after allowing for recombination and/or mixed infections. We refer to this as the PLA1 lineage. The remaining two samples with the amplification were from northwest Thailand and had a markedly different haplotype.

**Figure 2.**
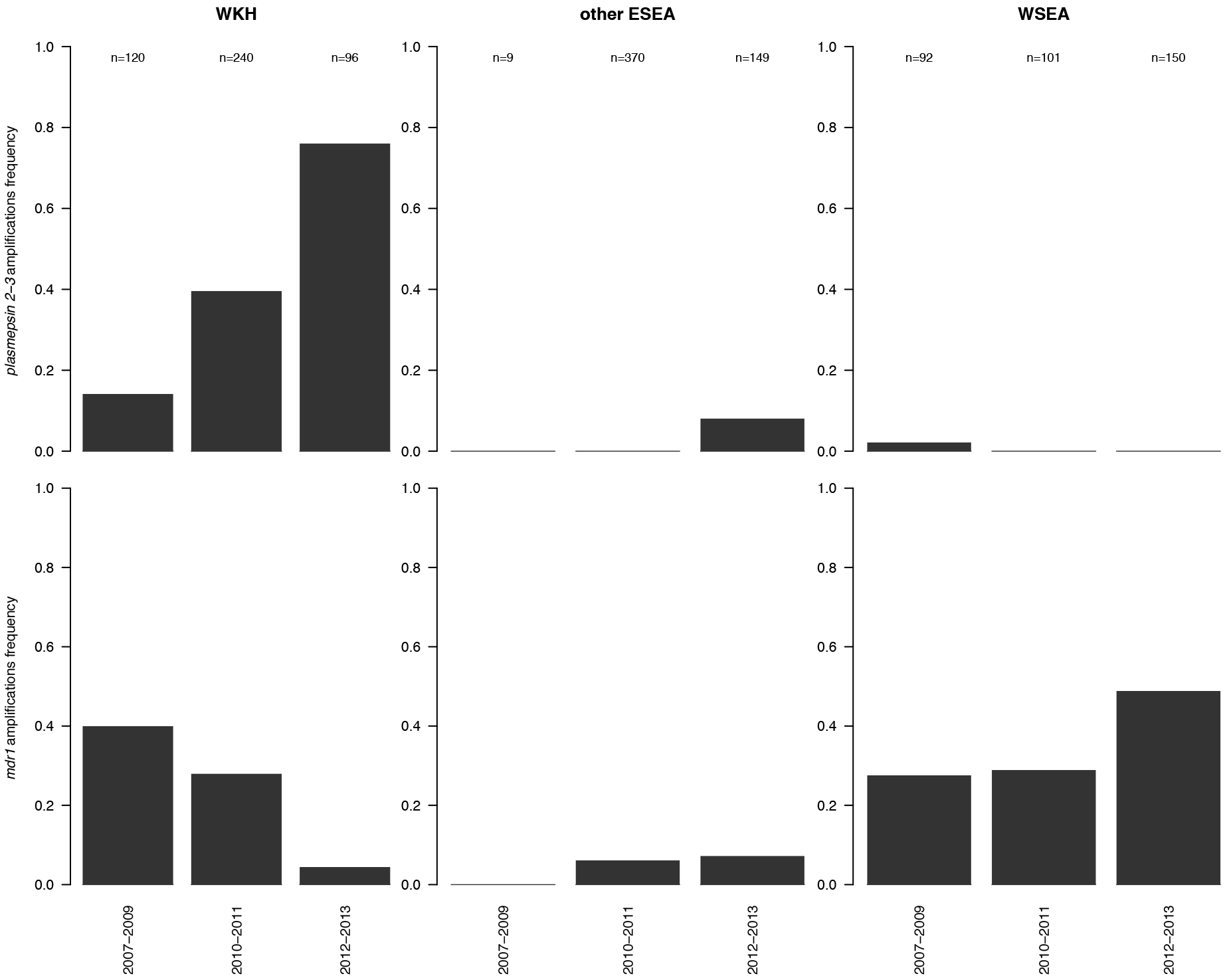
Frequency of *plasmepsin 2-3* (top) and mdrl (bottom) amplifications. The panels report the frequency of *plasmepsin 2-3* (top) and *mdrl* (bottom) amplifications in three consecutive year groups in western Cambodia (left panels); northern and northeastern Cambodia, Vietnam, Laos, and northeastern Thailand (central panels); and northwestern and southern Thailand and Myanmar (right panels). The sample size for each group is reported at the top.

In western Cambodia, *plasmepsin 2-3* amplifications were absent in 2007 but then rapidly increased to 78% frequency by 2013. This is in sharp contrast with the Thailand-Myanmar border, where only two amplifications were observed in 2008 and none after that. In northern Cambodia *plasmepsin 2-3* amplifications were first observed in 11 samples collected in 2012 and 2013.

### Emergence of DHA-PPQ resistance

The combination of a *kelch13* resistance allele with *plasmepsin 2-3* amplification is a marker of DHA-PPQ treatment failure.^10^ By analysing this combination of markers, we identified 154 samples as DHA-PPQ-resistant, after excluding samples with heterozygous genotypes, which were unsuitable for this analysis. A notable finding was that 94% (145/154) of DHA-PPQ-resistant samples carried the *kelch13* 580Y resistance allele, and 91% (140/154) belonged to the KEL1 lineage. A further 4% (6/154) of DHA-PPQ-resistant samples carried the 493H resistance allele and belonged to the KEL2 lineage (Supplementary Table 4). We identified 15 samples from western Cambodia and two from Thailand that carried *plasmepsin 2-3* amplifications but lacked a *kelch13* resistance allele.

These data indicate that DHA-PPQ resistance was present in western Cambodia as early as 2008, and then rapidly increased in frequency, from 16% in 2008 to 68% in 2013 (Figure 3). From the time of its emergence it was linked to the KEL1 lineage, which was observed in 80% (8/10) of DHA-PPQ-resistant samples in western Cambodia in 2008 and 93% (55/59) in 2012-2013. Over the same period there was an expansion of the proportion of KEL1 samples carrying *plasmepsin 2-3* amplifications (all of the PLA1 lineage) which increased from 31% (4/13) in 2008 to 92% (22/24) in 2013.

**Figure 3.**
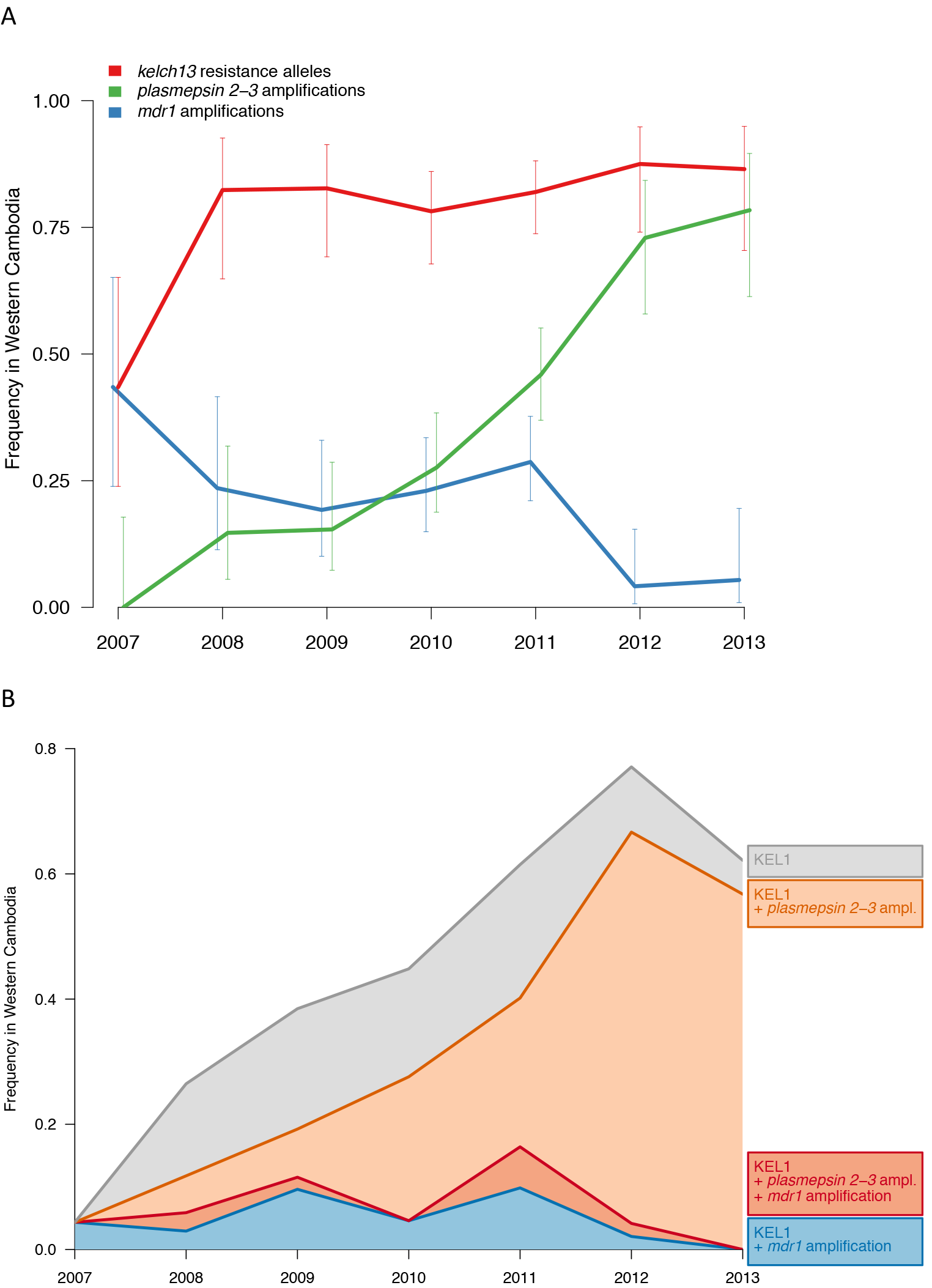
Frequency in western Cambodia of molecular markers for the three most commonly used drugs in the area and of the KEL1 lineage from 2007 to 2013. (A) The graph reports the cumulative frequency over time of *kelch13* resistance alleles (red), *plasmepsin 2-3* amplifications (green), and *mdr1* amplifications (blue). Error bars report the 95% confidence interval of the frequency, based on the sample size. (B) Frequency over time in western Cambodia of the dominant haplogroup KEL1. The coloured part, altogether, reports the annual prevalence of KEL1. Samples carrying the KEL1 haplogroup are divided into four groups according to the other markers (bottom to top): parasites with *mdr1* amplifications only (blue), parasites with *plasmepsin 2-3* and *mdrl* amplifications (red), parasites with *plasmepsin 2-3* amplifications only (orange), and parasites with single copies of both *plasmepsin 2-3* and *mdrl* (grey).

The acquisition of *plasmepsin 2-3* amplifications by parasites carrying the *kelch13* 493H resistance allele appears to be more recent, and this accounts for six of the DHA-PPQ resistant samples observed in western Cambodia between 2010 and 2013 (Supplementary Table 4). Taken together, the genomic evidence suggests that this acquisition occurred through introgression via recombination, possibly with parasites belonging to the KEL1 lineage.

### Geographic spread of DHA-PPQ resistance

Based on these genetic data, DHA-PPQ resistance appears to have been confined to western Cambodia until 2011, but in 2012-2013 it was observed in 11 samples from northern Cambodia and one sample from Laos close to the northern Cambodian border. Analysis of the *kelch13* and *plasmepsin 2-3* loci in these northern Cambodian samples showed that they resembled the majority of DHA-PPQ-resistant samples from western Cambodia, i.e. they all belonged to the KEL1/PLA1 colineage. This raises the strong possibility that DHA-PPQ resistance had spread from western to northern Cambodia.

To examine this question in more detail, we performed statistical chromosome painting across the whole genome and estimated co-ancestry between samples from different geographical locations. This revealed that, in northern Cambodia, samples with markers of DHA-PPQ resistance had much higher levels of western Cambodian co-ancestry than the rest of the population (Figure 4A). Construction of a neighbour-joining tree using whole genome data showed that northern Cambodian samples with markers of DHA-PPQ resistance grouped closely with parasites from western Cambodian part of the tree, rather than with other parasites from northern Cambodian (Figure 4B). Taken together, these findings indicate that the emergence of DHA-PPQ resistance in northern Cambodia in 2012 was the result of the migration of parasites from western Cambodia.

**Figure 4.**
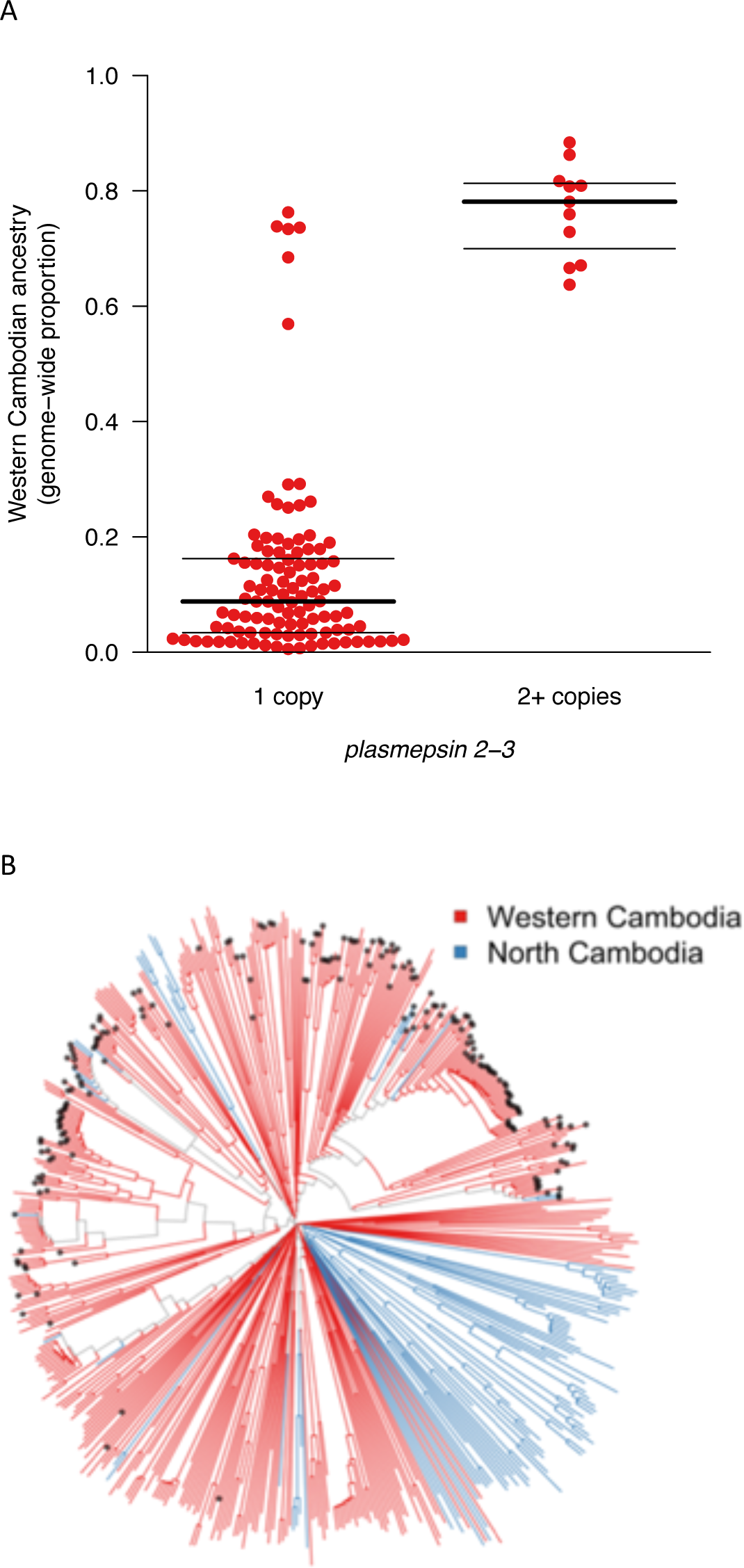
Spread of DHA-PPQ resistance to North Cambodia. (A) Each point represents a sample from northern Cambodia and represent the genome-wide proportion of western Cambodian ancestry for parasites with single (left) or multiple (right) copies of *plasmepsin 2-3*. Bold and thin horizontal lines indicate the median and IQR of each distribution, respectively. (B) Genome-wide neighbour-joining tree of all northern (blue) and western (red) Cambodian samples in the dataset, with those carrying *plasmepsin 2-3* amplifications identified by black dots at the tip.

### Declining frequency of *mdr1* amplifications

Artesunate-mefloquine was the first-line ACT in Cambodia prior to the adoption of DHA-PPQ in 2008, and has continued to be widely used in other parts of Southeast Asia.^20^ It has previously been noted that increasing piperaquine resistance in western Cambodia was accompanied by a reduction in mefloquine resistance.^13^

We further explored this relationship by analysing *mdr1* amplifications that are known markers of mefloquine resistance.^21^ Between 2007 and 2013, the frequency of *mdr1* amplifications in western Cambodia fell from 59% to 6%, in marked contrast to Myanmar and Thailand where the frequency rose from 13% in 2007 to 63% in 2013. This local decline in *mdr1* amplifications is directly related to the evolution and expansion of parasites belonging to the KEL1 haplogroup (Figure 3). In 2007, when the KEL1 lineage was at a frequency of 10% in western Cambodia, all KEL1 samples had *mdr1* amplifications and none had *plasmepsin 2-3* amplifications. Over the next few years, as the KEL1 lineage rose in frequency it progressively lost *mdr1* amplifications and gained *plasmepsin 2-3* amplifications. By 2012, when the KEL1 lineage had reached a frequency of 75% in western Cambodia, 92% of the KEL1 samples had *plasmepsin 2-3* amplifications whereas none had *mdr1* amplifications.

## Discussion

There has been much debate about whether the rapid spread of DHA-PPQ resistance in Southeast Asia presents a significant risk of derailing malaria control worldwide. ^14,16,22,23^. The World Health Organisation has determined that it is not a public health emergency, but this view is challenged by some malaria experts.^22,23^ Here we report a longitudinal genomic analysis of the *P. falciparum* population of western Cambodia, over a 7 year period encompassing the emergence and expansion of this outbreak of multidrug-resistant malaria. The findings shed light on four questions that are germane to the ongoing debate: (i) what is the nature of the parasites that are causing the problem; (ii) how rapidly have they spread; (iii) how are they likely to respond to alternative drug combinations; (iv) what are the major risks and uncertainties in the longer term?

First, in population genetic terms, what are the essential features of DHA-PPQ-resistant parasites? In the present study, 91% carried a *kelch13* 580Y allele of the KEL1 lineage, and 4% carried a *kelch13* 493H allele of the KEL2 lineage. Thus most but not all DHA-PPQ resistant parasites acquired artemisinin resistance from a single epidemiological origin. This is a notable finding, since we observed 38 putative independent origins of artemisinin resistance.

At the *plasmepsin 2-3* locus we identified only two epidemiological origins of gene amplifications associated with piperaquine resistance. One of these was relatively rare, being found in only two parasites from Thailand. All of the DHA-PPQ-resistant parasites belonged to a single *plasmepsin 2-3* lineage that we refer to as PLA1.

These findings are concordant with recent reports of DHA-PPQ-resistant parasites found in Thailand, Laos and Vietnam.^14,16^ Imwong and colleagues refer to this as the *PfPailin* lineage, which might be taken to mean that this is a uniform type of *P. falciparum*, analogous to a viral or bacterial strain. However malaria parasites, unlike viruses and bacteria, undergo sexual recombination with each transmission cycle therefore this nomenclature could potentially be misleading. These data show that the KEL1 lineage of artemisinin resistance is now carried by parasites of diverse genetic backgrounds (Supplementary Figure 2) and, while parasites of the KEL1/PLA1 co-lineage have high levels of shared ancestry relative to the general parasite population, they are not a genetically homogeneous entity. It is also important to note that the KEL1 and PLA1 lineages are separable and possibly emerged independently, but here they have come together to form a co-lineage that we refer to as KEL1/PLA1. While this might seem a technical detail, it is central to understanding how this form of multidrug resistance emerged, and how it might evolve in future by recombining with other parasites and incorporating new genetic features.

Second, when did this group of parasites emerge and how rapidly have they spread? These data show that parasites with KEL1 lineage were present at low frequency in western Cambodia as early as 2007. At that time, they carried *mdr1* amplifications that confer mefloquine resistance and they did not carry *plasmepsin 2-3* amplifications. The KEL1/PLA1 co-lineage was first observed in 2008, the same year that DHA-PPQ was introduced as the first-line antimalarial drug in this part of the country. It is possible that PLA1 existed beforehand at low frequency, independent of KEL1, since piperaquine was previously used as a monotherapy in Cambodia. Over the time period and in the locations surveyed here, PLA1 has been mainly linked to KEL1, but it is also seen in parasites with the KEL2 haplogroup and in parasites without *kelch13* resistance alleles.

Prior to the emergence of DHA-PPQ resistance there was already a very high frequency of artemisinin resistant parasites in western Cambodia, but they comprised a diverse set of *kelchi* resistance alleles arising from multiple epidemiological origins, each of which tended to remain relatively localised. After the KEL1/PLA1 co-lineage emerged in 2008 it spread rapidly and extensively across western Cambodia, reaching a frequency of over 60% in the parasite population. These data show that it appeared in northern Cambodia in 2012, and that this was due to spread of parasites from western Cambodia. Although the genetic typing methods used by Imwong and colleagues to characterise *kelch13* haplotypes and *plasmepsin 2-3* amplifications are different from those used here, there seems little doubt from their data that the DHA-PPQ-resistant parasites that appeared in northeastern Thailand and Laos in 2014-5, and in Vietnam in 2016, correspond to the KEL1/PLA1 co-lineage.^14,16^

There are many parallels between drug resistance outbreaks and cancer. In cancer diagnosis, the term ‘aggressive’ means a tumour that is rapidly spreading, as opposed to one that remains localised. By this definition, the KEL1/PLA1 co-lineage can be described as an aggressive form of drug resistance.

Third, what short term predictions can be made about the likely response of DHA-PPQ-resistant parasites to alternative drug combinations? As noted above, the KEL1 lineage was initially associated with *mdr1* amplifications that are markers of mefloquine resistance, but this association has broken down over time, and the most recent KEL1/PLA1 samples in this dataset do not carry *mdr1* amplifications. Therefore DHA-PPQ-resistant parasites are currently sensitive to mefloquine, and artesunate-mefloquine is now being used successfully as first line antimalarial treatment in Cambodia. There is also evidence that DHA-PPQ resistant parasites are responsive to artesunate-pyronaridine, a new form of ACT.^22^

It is reasonable to expect that the switch to artesunate-mefloquine in Cambodia will cause levels of piperaquine resistance to fall, but that it will also cause levels of mefloquine resistance to rise. KEL1 parasites in particular have evidently no problem in switching between mefloquine and piperaquine resistance: in 2007 they all carried *mdr1* amplifications and none carried *plasmepsin 2-3* amplifications, whereas by 2013 this was almost completely reversed. However, there is evidence of natural antagonism between piperaquine resistance and mefloquine resistance, i.e. piperaquine usage may naturally act to drive down levels of mefloquine resistance and vice versa (refs ^10,11^; Rutledge et al, unpublished data). This raises the possibility that the effectiveness of ACT in Southeast Asia could be maintained by using mefloquine and piperaquine in combination or in rotation, but any such strategy will need to be closely monitored for the emergence of joint resistance to both drugs, which needs to be avoided at all costs.

Finally, what are the longer-term risks and how might it be possible to reduce the level of uncertainty? The main risks are that *P. falciparum* malaria will eventually become untreatable in Southeast Asia, and that this will spread to Africa which would be catastrophic. The aggressive spread of KEL1/PLA1 suggests that artemisinin resistant parasites are acquiring a higher level of biological fitness, and the question is to what extent that increases the risk of partner drug failure and of trans-continental spread. Some might argue that the risk is relatively low, e.g. the success of KEL1 lineage might have been simply due to its association with PLA1 at a time when DHA-PPQ was the first-line antimalarial, in which case it can potentially be contained by strategic management of DHA-PPQ usage. On the other hand, it can be argued that KEL1 appears to have aggressive characteristics that are independent of its association with PLA1. It carries the 580Y allele, which has more independent origins than any other artemisinin resistance allele (Supplementary Table 2). The 580Y allele has successfully outcompeted other artemisinin resistance alleles, not only in western Cambodia but also on the Thailand-Myanmar border, ^24^ and since it has emerged independently in these two locations, this superior fitness would appear to be independent of its genetic background. It is also possible that the KEL1 lineage is progressively undergoing evolutionary adaptation by becoming linked to compensatory or synergistic mutations in other genes that act to increase its transmissibility and biological fitness.

Tools to monitor evolutionary changes in the parasite population would enable national malaria control programmes to know if and when interventions are going wrong, reduce the level of uncertainty and, ultimately, minimise the risks associated. In retrospect, we can see that the KEL1/PLA1 co-lineage emerged within the first year of use of DHA-PPQ as a first line antimalarial in western Cambodia, and had risen to a frequency of 60% by the time that the first clinical reports of the problem emerged five years later. Prospective genomic surveillance is now called for, and not simply for known resistance markers, as the major risks and uncertainties pertain to future evolutionary events. With appropriate technologies and concerted action, it ought to be possible to avoid similar situations arising in the future.

## Methods

### Data

This publication uses data from a clinical study of DHA-PPQ failure in Cambodia.^25^ To contextualise the findings, they were complemented with open data generated by the MalariaGEN Plasmodium *falciparum* Community Project.^17^ For all the samples analysed, sequence data were generated at the Wellcome Trust Sanger Institute using Illumina short-read technology and genotypes were called with a standardised analysis pipeline.^26^

We only utilised samples from south/southeast Asia with sufficient coverage (more than 75% of the SNP covered by at least 5 reads) and with low complexity of infection (*F*_*WS*_ > 0.8). This is to minimise the risk that the multi-locus analyses and haplotype analyses are confounded by complex infections.

### *Kelch13* genotyping and haplogroup assignment

The genotype of *kelch13* per each sample was derived from read counts at non-synonymous SNPs in the propeller and in the BTB/POZ domains using a procedure described previously.^7^

We reconstructed the probable origin of *kelch13* mutation using chromosome painting.^19^ This method compares haplotypes in a sample to those in the remaining samples, and estimates the probability that a genome fragment originates in each of them while also accounting for recombination and *de novo* mutations.

For all and only *kelch 13* homozygous mutant samples (n=553), we ran chromosome painting on the entire chromosome 13, obtaining posterior copying probabilities for all loci (an approximation of the probability of two samples being closest neighbours, for each locus, in the underlying genealogy). Mutation rate (i.e. miscopying parameter) was estimated per sample using the Watterson’s estimator, and we assumed a uniform recombination map with a rate of 500 kbp/cM. The scaling parameter (termed effective population size) was set to 1000. To account for the presence of residual heterozygous genotypes due to mixed infections, ε (the probability of emitting a mixed call) was set to 10^−8^.^17^ We repeated the analysis varying this parameter set, to assess the effects of misspecification, and results were found to be very similar qualitatively (data not shown). We aggregated these probabilities by taking, for each sample, their average inside the boundaries of the *kelch13* gene (Pf3D7_13_v3: 1724817–1726997). Different aggregation methods, including utilising single variants inside the gene, produced identical results (data not shown). This process resulted in one “copying vector” *k*_*i*_ of 553 elements per sample *i* (*i* = 1,…,553) reporting the probability of that sample being close to all remaining ones in the underlying genealogy of the *kelch13* locus.

To assign samples to haplogroups, these copying vectors *k*_*i*_ were assembled together to form a matrix *K* of dimension 553×553, which can be interpreted as a measure of ancestral similarity between all pairs of samples. We performed a complete hierarchical clustering using as distance matrix the complement of log(*K* + *K*^*T*^) (Supplementary Figure 4A). This analysis was performed using the function `hclust(method=“complete”)` as implemented in R version 3.3.2.^27^ The resulting dendrogram was subsequently cut to produce discreet clusters. The cut-off was determined heuristically in order to maximise cluster homogeneity by visually inspecting: (i) the haplotype structure (Supplementary Figure 1); (ii) the distance matrix (Supplementary Figure 4A); and (iii) a t-SNE reduction of the distance matrix (Supplementary Figure 4B). This process identified 24 distinct clusters. The t-SNE representation was calculated using the R function and package tsne and ran with default parameters.^28^ Finally, haplogroups were defined as groups of samples having the same *kelch13* resistance allele and belonging to the same cluster.

### Copy number alleles

We used two orthogonal methods to determine duplication genotypes around *mdr1* and *plasmepsin 2-3*: a coverage-based method and a method based on position and orientation of reads near discovered duplication breakpoints. Details of the methods are described elsewhere (MalariaGEN *Plasmodium falciparum* Community Project, 2017, manuscript in preparation). In short, a coverage based hidden Markov model (HMM) was firstly used to identify potential copy number amplifications and their boundaries.^29^ Following this we manually determined breakpoints of duplications around *mdr1* and *plasmepsin 2-3* by visual inspection of soft clipped reads and paired reads aligning either facing away from each other or aligned in the same orientation. We then searched all samples for breakpoint-spanning reads, and combined results with those of the HMM.

### Spread of DHA-PPQ resistance

We estimated the level of co-ancestry between different locations by using chromosome painting. The method was run using the same parameters specified above, but across the whole genome. For each sample, we generated the most likely painting (i.e. Viterbi decoding) and aggregated copying vectors according to the geographical origin of the donor samples. We then estimated the fraction of the genome copied from each population.

The neighbour-joining trees were produced using the function `nj` in the R package `ape`
, with default parameters and using a genome-wide pairwise genetic distance matrix as described elsewhere.^17,30^

## Contributors

RA, OM, RMF, and DPK designed the study. CA, PL, SSu, SSr and ED collected the samples and/or carried out laboratory work to sequence the samples. RA, RDP, JAG, JWS, and OM generated and curated the data. RA, RDP, JAG, OM and DPK analysed and interpreted the data. RA and DPK wrote the manuscript.

## Acknowledgments

This work was funded by the Bill & Melinda Gates Foundation (OPP1040463, OPP11188166); Medical Research Council (G0600718); UK Department for International Development (M006212); and Intramural Research Program of the National Institute of Allergy and Infectious Diseases, National Institutes of Health. This publication uses data from the MalariaGEN *Plasmodium falciparum* Community Project. Genome sequencing was done by the Wellcome Trust Sanger Institute with funding from the Wellcome Trust (098051, 206194). The Community Project was coordinated by the MalariaGEN Resource Centre with funding from the Wellcome Trust (090770, 204911). We thank the staff of the WTSI Sample Logistics, Sequencing, and Informatics facilities for their contribution; all patients and collaborators contributing samples to the MalariaGEN *Plasmodium falciparum* Community Project; Mihir Kekre and Katie Love for their support in the sample processing pipeline; and Sonia Goncalves, Victoria Cornelius and Christa Henrichs for their support in coordinating and managing the project and the MalariaGEN *Plasmodium falciparum* Community Project.

## Supplementary material

**Supplementary Table 1.** Number of *kelch13* wild-type, heterozygous, and mutant samples in the dataset. SAS = South Asia; WSEA = Southeast Asia - West; ESEA = = Southeast Asia - East.

| *kelch13* |  | SAS | WSEA |  | ESEA |  |  |  |  |  |  |
| --- | --- | --- | --- | --- | --- | --- | --- | --- | --- | --- | --- |
| *kelch13* | *kelch13* resistance allele | Bangladesh | Myanmar | Thailand (Northwest and South) | Cambodia (North) | Cambodia (Northeast) | Cambodia (West) | Vietnam | Thailand (Northeast) | Laos | Total |
| Wild-type |  | 53 | 60 | 207 | 86 | 121 | 80 | 103 | 1 | 92 | 803 |
| Het |  | 1 | 7 | 40 | 4 | 1 | 61 | 17 | 3 | 2 | 136 |
| Mutant | Total |  | 34 | 86 | 33 | 3 | 323 | 57 | 15 | 2 | 553 |
|  | 580Y |  | 11 | 24 | 26 | 3 | 241 | 9 | 3 |  | 317 |
|  | 493H |  |  |  | 5 |  | 47 | 4 |  |  | 56 |
|  | 539T |  |  |  | 2 |  | 23 | 4 | 12 | 2 | 43 |
|  | 543T |  |  |  |  |  | 2 | 22 |  |  | 24 |
|  | 441L |  | 5 | 11 |  |  |  |  |  |  | 16 |
|  | 561H |  | 2 | 13 |  |  |  |  |  |  | 15 |
|  | 675V |  | 2 | 13 |  |  |  |  |  |  | 15 |
|  | 553L |  |  | 2 |  |  |  | 9 |  |  | 11 |
|  | 538V |  |  | 8 |  |  |  |  |  |  | 8 |
|  | 449A |  | 2 | 3 |  |  | 2 |  |  |  | 7 |
|  | 574L |  | 6 | 1 |  |  |  |  |  |  | 7 |
|  | 353Y |  |  |  |  |  |  | 5 |  |  | 5 |
|  | 527H |  |  | 5 |  |  |  |  |  |  | 5 |
|  | 568G |  |  |  |  |  |  | 4 |  |  | 4 |
|  | 481V |  |  | 2 |  |  | 2 |  |  |  | 4 |
|  | 446I |  | 3 |  |  |  |  |  |  |  | 3 |
|  | 719N |  |  |  |  |  | 3 |  |  |  | 3 |
|  | 673I |  | 2 |  |  |  |  |  |  |  | 2 |
|  | 584V |  |  |  |  |  | 2 |  |  |  | 2 |
|  | 438N |  | 1 | 1 |  |  |  |  |  |  | 2 |
|  | 443S |  |  | 1 |  |  |  |  |  |  | 1 |
|  | 395Y |  |  |  |  |  | 1 |  |  |  | 1 |
|  | 614L |  |  | 1 |  |  |  |  |  |  | 1 |
|  | 537I |  |  | 1 |  |  |  |  |  |  | 1 |

**Supplementary Table 2.** Geographical Distribution Of The 38 *Kelch13* Haplogroups.

| *kelch13* haplogroup | Samples | *kelch13* resistance allele | Cambodia (West) | Cambodia (North) | Cambodia (Northeast) | Laos | Thailand (Northeast) | Vietnam | Myanmar | Thailand (Northwest and South) |
| --- | --- | --- | --- | --- | --- | --- | --- | --- | --- | --- |
| KEL1 | 266 | 580Y | 226 | 26 | 2 |  | 2 | 9 |  | 1 |
| KEL2 | 49 | 493H | 40 | 5 |  |  |  | 4 |  |  |
| KEL3 | 40 | 539T | 21 | 2 |  | 2 | 11 | 4 |  |  |
| KEL4 | 28 | 580Y |  |  | 1 |  | 1 |  | 11 | 15 |
| KEL5 | 24 | 543T | 2 |  |  |  |  | 22 |  |  |
| KEL6 | 15 | 675V |  |  |  |  |  |  | 2 | 13 |
| KEL7 | 13 | 580Y | 13 |  |  |  |  |  |  |  |
| KEL8 | 11 | 561H |  |  |  |  |  |  |  | 11 |
| KEL9 | 9 | 553L |  |  |  |  |  | 7 |  | 2 |
| KEL10 | 8 | 580Y |  |  |  |  |  |  |  | 8 |
| KEL11 | 8 | 538V |  |  |  |  |  |  |  | 8 |
| KEL12 | 8 | 441L |  |  |  |  |  |  |  | 8 |
| KEL13 | 7 | 574L |  |  |  |  |  |  | 6 | 1 |
| KEL14 | 7 | 449A | 2 |  |  |  |  |  | 2 | 3 |
| KEL15 | 5 | 441L |  |  |  |  |  |  | 3 | 2 |
| KEL16 | 5 | 527H |  |  |  |  |  |  |  | 5 |
| KEL17 | 4 | 493H | 4 |  |  |  |  |  |  |  |
| KEL18 | 4 | 568G |  |  |  |  |  | 4 |  |  |
| KEL19 | 4 | 353Y |  |  |  |  |  | 4 |  |  |
| KEL20 | 4 | 561H |  |  |  |  |  |  | 2 | 2 |
| KEL21 | 4 | 481V | 2 |  |  |  |  |  |  | 2 |
| KEL22 | 3 | 493H | 3 |  |  |  |  |  |  |  |
| KEL23 | 3 | 441L |  |  |  |  |  |  | 2 | 1 |
| KEL24 | 3 | 719N | 3 |  |  |  |  |  |  |  |
| KEL25 | 3 | 446I |  |  |  |  |  |  | 3 |  |
| KEL26 | 2 | 553L |  |  |  |  |  | 2 |  |  |
| KEL27 | 2 | 539T | 1 |  |  |  | 1 |  |  |  |
| KEL28 | 2 | 438N |  |  |  |  |  |  | 1 | 1 |
| KEL29 | 2 | 584V | 2 |  |  |  |  |  |  |  |
| KEL30 | 2 | 673I |  |  |  |  |  |  | 2 |  |
| KEL31 | 1 | 580Y | 1 |  |  |  |  |  |  |  |

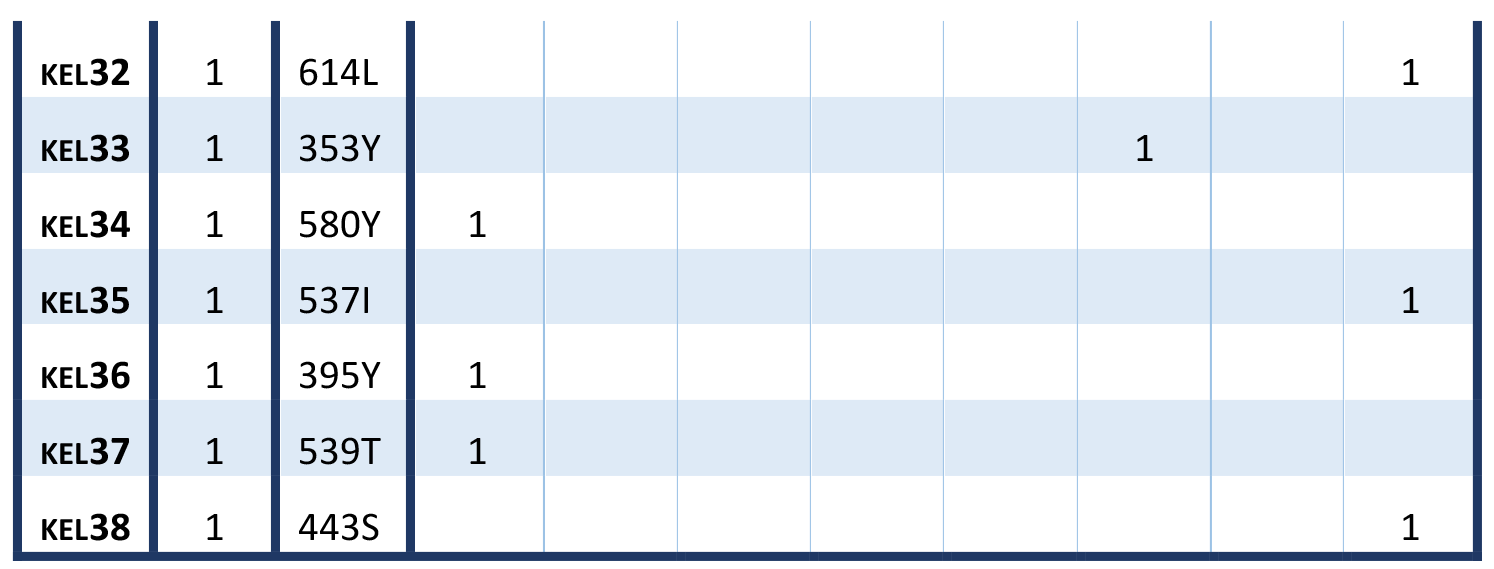

**Supplementary Table 3.** Temporal and geographical distribution of samples carrying *plasmepsin* 2-3 amplication.

|  | Cambodia (West) | Cambodia (North) | Cambodia (Northeast) | Thailand (Northwest) |
| --- | --- | --- | --- | --- |
| 2008 | 8 |  |  | 2 |
| 2009 | 9 |  |  |  |
| 2010 | 31 |  |  |  |
| 2011 | 64 |  |  |  |
| 2012 | 43 | 4 |  |  |
| 2013 | 30 | 7 | 1 |  |
| **Total** | **185** | **11** | **1** | **2** |

**Supplementary Table 4.**
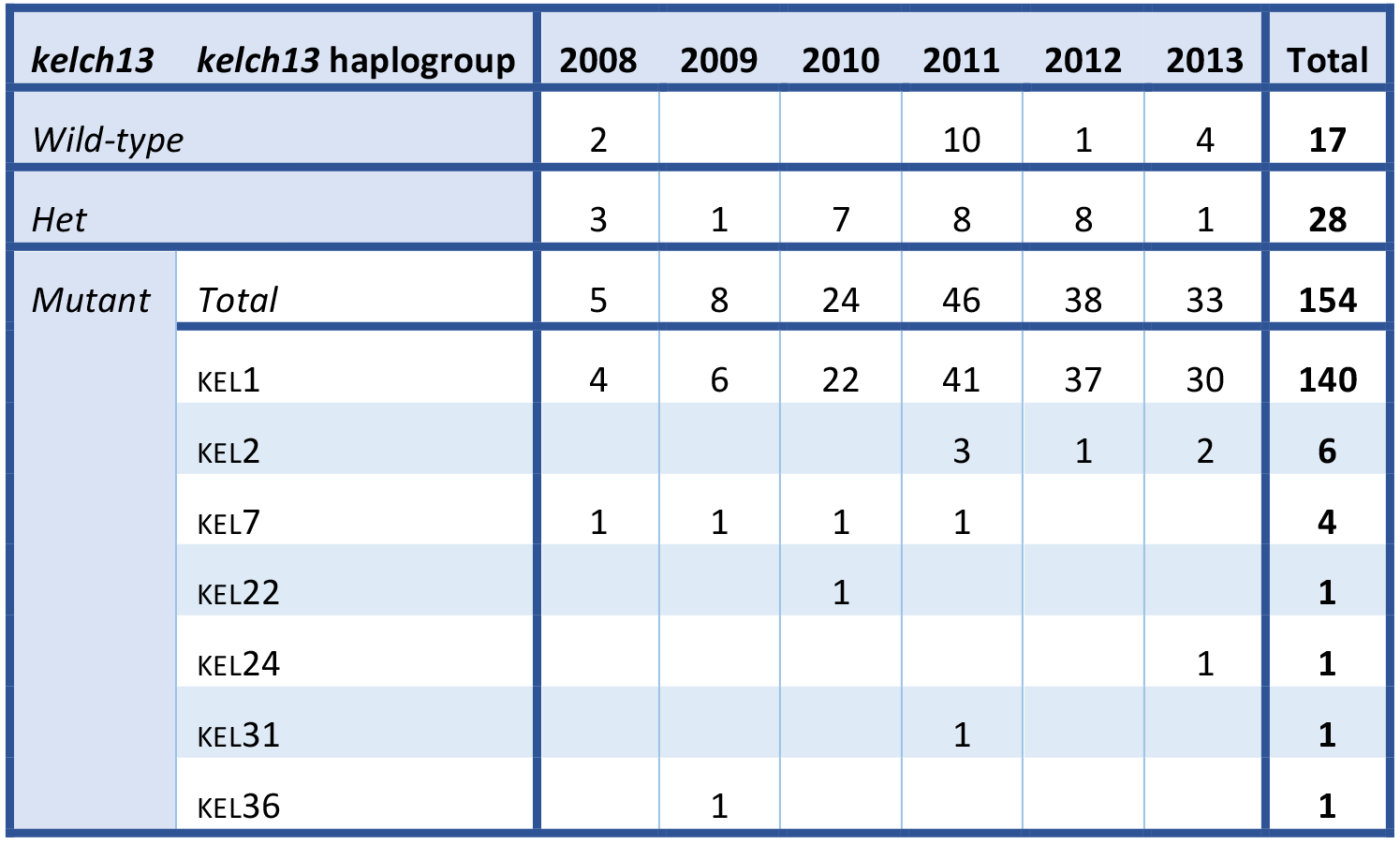
Distribution of samples carrying *plasmepsin 2-3* amplification across the *kelch13* haplogroups.

| *kelch13* | *kelch13* haplogroup | 2008 | 2009 | 2010 | 2011 | 2012 | 2013 | Total |
| --- | --- | --- | --- | --- | --- | --- | --- | --- |
| *Wild-type* |  | 2 |  |  | 10 | 1 | 4 | **17** |
| *Het* |  | 3 | 1 | 7 | 8 | 8 | 1 | **28** |
| *Mutant* | *Total* | 5 | 8 | 24 | 46 | 38 | 33 | **154** |
|  | KEL1 | 4 | 6 | 22 | 41 | 37 | 30 | **140** |
|  | KEL2 |  |  |  | 3 | 1 | 2 | **6** |
|  | KEL7 | 1 | 1 | 1 | 1 |  |  | **4** |
|  | KEL22 |  |  | 1 |  |  |  | **1** |
|  | KEL24 |  |  |  |  |  | 1 | **1** |
|  | KEL31 |  |  |  | 1 |  |  | **1** |
|  | KEL36 |  | 1 |  |  |  |  | **1** |

**Supplementary Figure 1.**
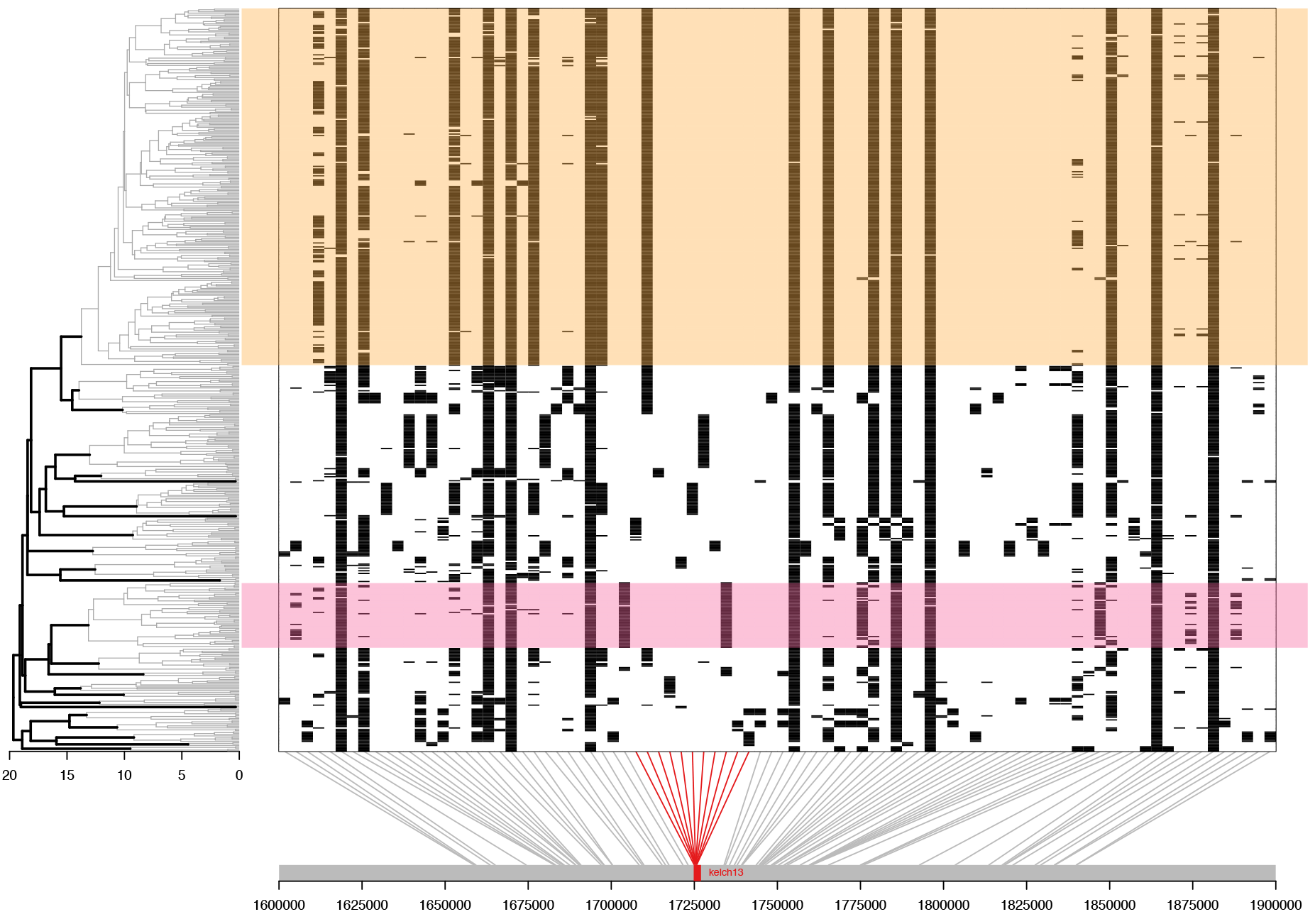
Haplotype analysis of samples carrying *kelch13* resistance alleles. Haplotype diagram where each vertical column represents a SNP (MAF >1%) within approximately 100 kbp either side of the *kelch13* gene, and each horizontal line a sample; at the intersection, black lines represents a non-reference genotype allele in the sample (a read majority call is used for heterozygous genotypes). Grey line at the bottom reports the position of the SNPs within chromosome 13, with the *kelch13* genes highlighted in red. A complete hierarchical clustering dendrogram is reported on the left, based on the co-ancestry matrix calculated using statistical chromosome painting. Bold black lines join clusters whose distance is above the cut-off point (height=14) while thin grey lines show the internal structure of each cluster. Highlighted in orange and red are samples belonging to KEL1 and KEL2, respectively.

**Supplementary Figure 2.**
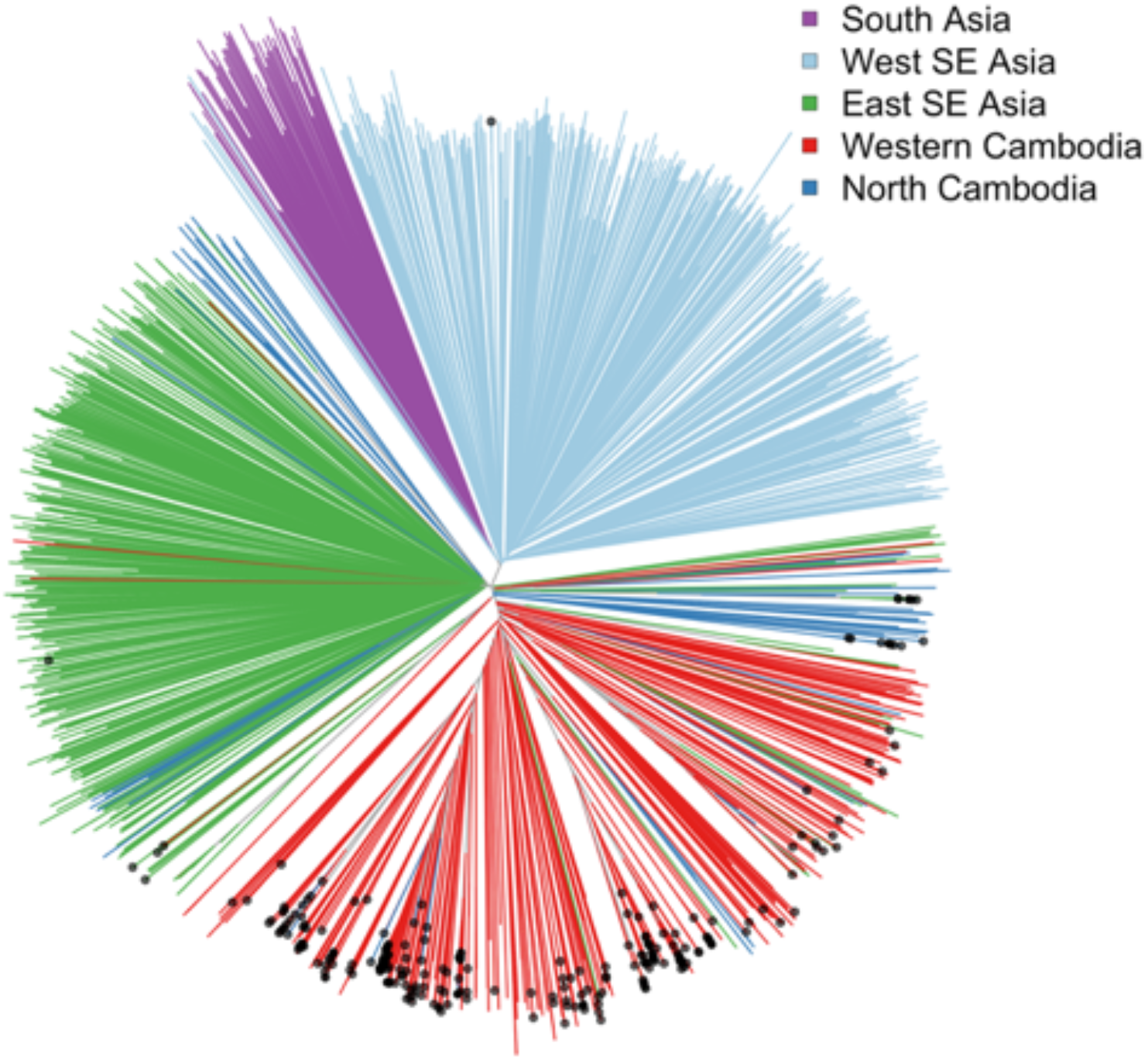
Genetic relatedness of KEL1 samples to all other samples. Genome-wide neighbour-joining tree of all samples in the dataset based on their overall genetic similarity. Each segment represents one sample and is coloured according to the region of collection. Parasites carrying the dominant haplogroup KEL1 are identified by a black dot at the tip.

**Supplementary Figure 3.**
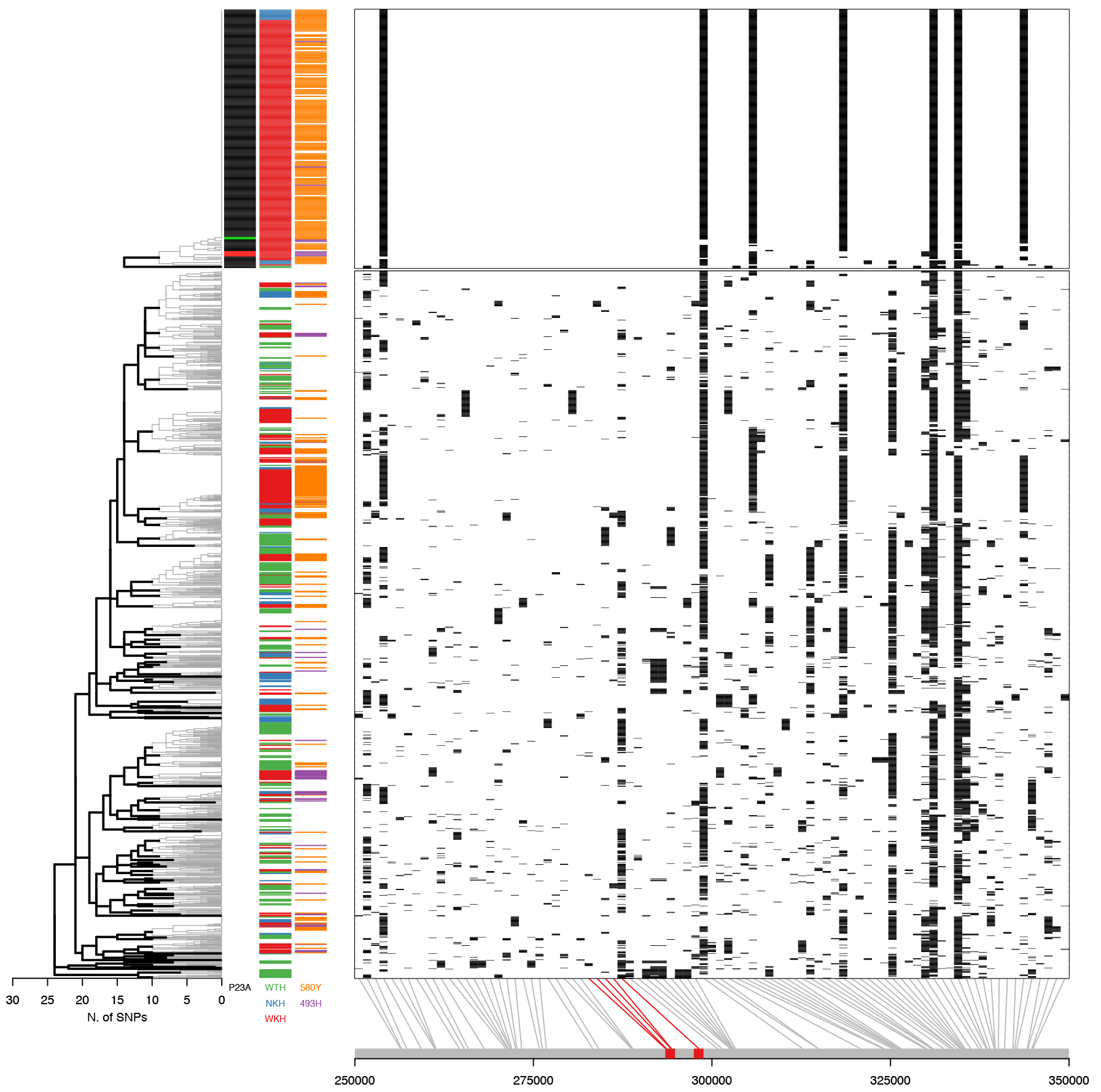
Analyses of haplotypes surrounding the *plasmepsin 2-3* genes. Haplotype diagram where each vertical column represents a SNP (MAF >1%) within 100 kbp around the *plasmepsin 2-3* genes, and each horizontal line a sample; at the intersection, black lines represents a non-reference genotype allele in the sample (a read majority call is used for heterozygous genotypes). Haplotypes are shown for samples carrying the amplification (top, n=199) and wild-type alleles (bottom, n=1266); 26 unclassified samples are not shown. The first column of coloured lines on the left (P23A) reports the presence of the amplification and the set of breakpoints identified (black, red, or green for each one of the three sets of breakpoints, white for wild-type). The second column reports the origin of the sample for the three major regions where amplifications are observed (WKH = western Cambodia, red; NKH = northern Cambodia, blue; WTH = northwestern Thailand, green; white = elsewhere). The last column reports the two major *kelch13* resistance alleles associated with the amplification (580Y = orange; 493H = purple). Grey line at the bottom reports the position of the SNPs within chromosome 14, with the *plasmepsin 2-3* genes highlighted in red. A complete hierarchical clustering dendrogram is reported on the left, with mutant and wild-type samples clustered independently for clarity. The height of the joints in the dendrogram reports the maximum number of differences between any two haplotypes.

**Supplementary Figure 4.**
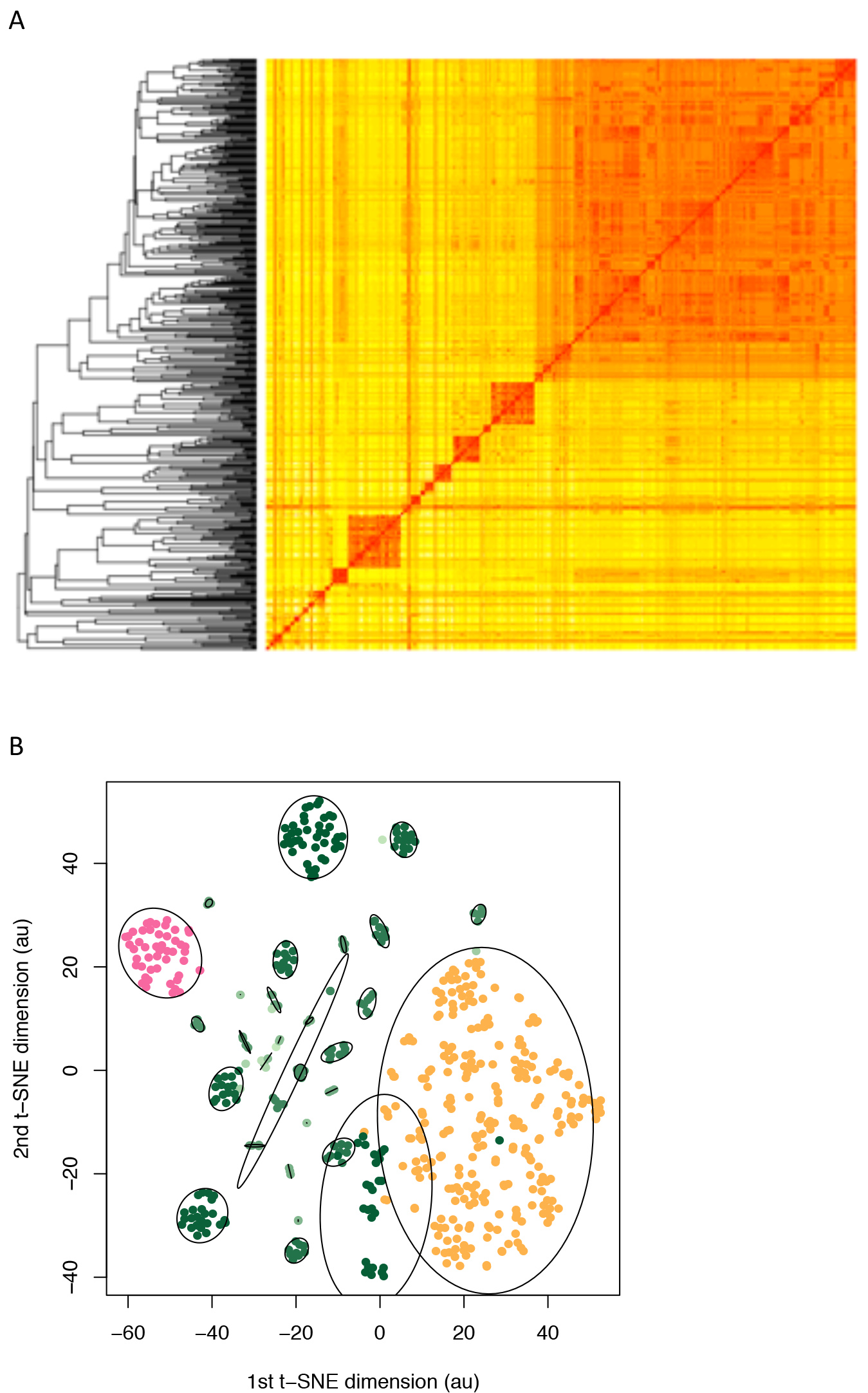
Co-ancestry matrix of 553 samples carrying homozygous *kelch13* resistance alleles. (A) The matrix has 553 rows and columns and is symmetric along the diagonal. The complete hierarchical clustering dendrogram is reported on the left. Red colours represent higher level of co-ancestry (arbitrary unit). (B) Two-dimensional visualization of the same coancestry matrix using t-SNE, a dimensionality reduction approach. Each dot represents a sample coloured according the to the *kelch13* haplogroup it belongs to, using the same colour scheme as Figure 1. Yellow and pink samples carry KEL1 and KEL2 haplogroups, respectively. Ellipses capture 90% of the spread of each haplogroup.

